# Systematics, biogeography and diversification of *Scytalopus* tapaculos (Rhinocryptidae), an enigmatic radiation of Neotropical montane birds

**DOI:** 10.1101/600775

**Authors:** Carlos Daniel Cadena, Andrés M. Cuervo, Laura N. Céspedes, Gustavo A. Bravo, Niels Krabbe, Thomas S. Schulenberg, Graham E. Derryberry, Luis Fabio Silveira, Elizabeth P. Derryberry, Robb T. Brumfield, Jon Fjeldså

## Abstract

We studied the phylogeny, biogeography and diversification of suboscine birds in the genus *Scytalopus* (Rhinocryptidae), a widespread, speciose, and taxonomically challenging group of Neotropical birds. We analyzed nuclear (exons, regions flanking ultraconserved elements) and mitochondrial (ND2) DNA sequence data for a taxonomically and geographically comprehensive sample of specimens collected from Costa Rica to Patagonia and eastern Brazil. We found that *Scytalopus* is a monophyletic group sister to *Eugralla*, and consists of three main clades roughly distributed in (1) the Southern Andes, (2) eastern Brazil, and (3) the Tropical Andes and Central America. The clades from the Southern Andes and Eastern Brazil are sister to each other. Despite their confusing overall uniformity in plumage coloration, body shape and overall appearance, rates of species accumulation through time in *Scytalopus* since the origin of the clade in the Late Miocene are unusually high compared to those of other birds, suggesting rapid non-adaptive diversification in the group which we attribute to their limited dispersal abilities making them speciation-prone and their occurrence in a complex landscape with numerous barriers promoting allopatric differentiation. Divergence times among species and downturns in species accumulation rates in recent times suggest that most speciation events in *Scytalopus* predate climatic oscillations of the Pleistocene. Our analyses identified various cases of strong genetic structure within species and lack of monophyly of taxa, flagging populations which likely merit additional study to establish their taxonomic status. In particular, detailed analyses of species limits are due in *S. parvirostris, S. latrans, S. speluncae*, the *S. atratus* complex, and the Southern Andes clade.

## Introduction

*Scytalopus* tapaculos (Rhinocryptidae) live in all major montane systems of South America and lower Central America except in the Pantepui, and extend into lowlands and foothills in southern South America and eastern Brazil (Figure 1a). Because these suboscine passerines have limited dispersal abilities and narrow elevational distributions (Krabbe and Schulenberg 2003; Cadena and Céspedes 2019) they are prone to track the historical dynamics of their habitats, which makes them an appropriate model to study the diversification of Neotropical montane organisms and to analyze the accumulation of species in mountains over time. Given the austral distribution of several Rhinocryptid genera representing deep branches in the family tree (Figure 1a), *Scytalopus* may have originated in southern South America and subsequently spread northwards as the Andes uplifted (Irestedt et al. 2002), but this hypothesis, a long-standing issue in the historical biogeography of Andean birds (Chapman 1917), has not been formally tested owing to the lack of a robust phylogeny. Likewise, hypotheses posed to account for the disjunct distribution of *Scytalopus* in eastern Brazil and in the Andes (Sick 1985; Vielliard 1990; Maurício 2005; Willis 1992) have not been amenable to testing. In addition to the lack of information on biogeographic history, little is known about the temporal pattern of diversification in *Scytalopus*. Based on the purportedly crucial role of Pleistocene climatic fluctuations on the speciation of Neotropical montane birds (Weir 2006; Vuilleumier 1969), one might predict high recent rates of diversification of the genus. Alternatively, considering that local diversity of *Scytalopus* is typically low (i.e., species are rarely syntopic, suggesting mutual exclusion through competitive interactions; Krabbe and Schulenberg 2003), rates of diversification might have been higher early on in the history of the group if mountain uplift promoted opportunities for isolation and diversification in new environments and declined towards recent times as a result of filling of ecological and geographical space (Rabosky 2009; Price et al. 2014).

**Figure 1.**
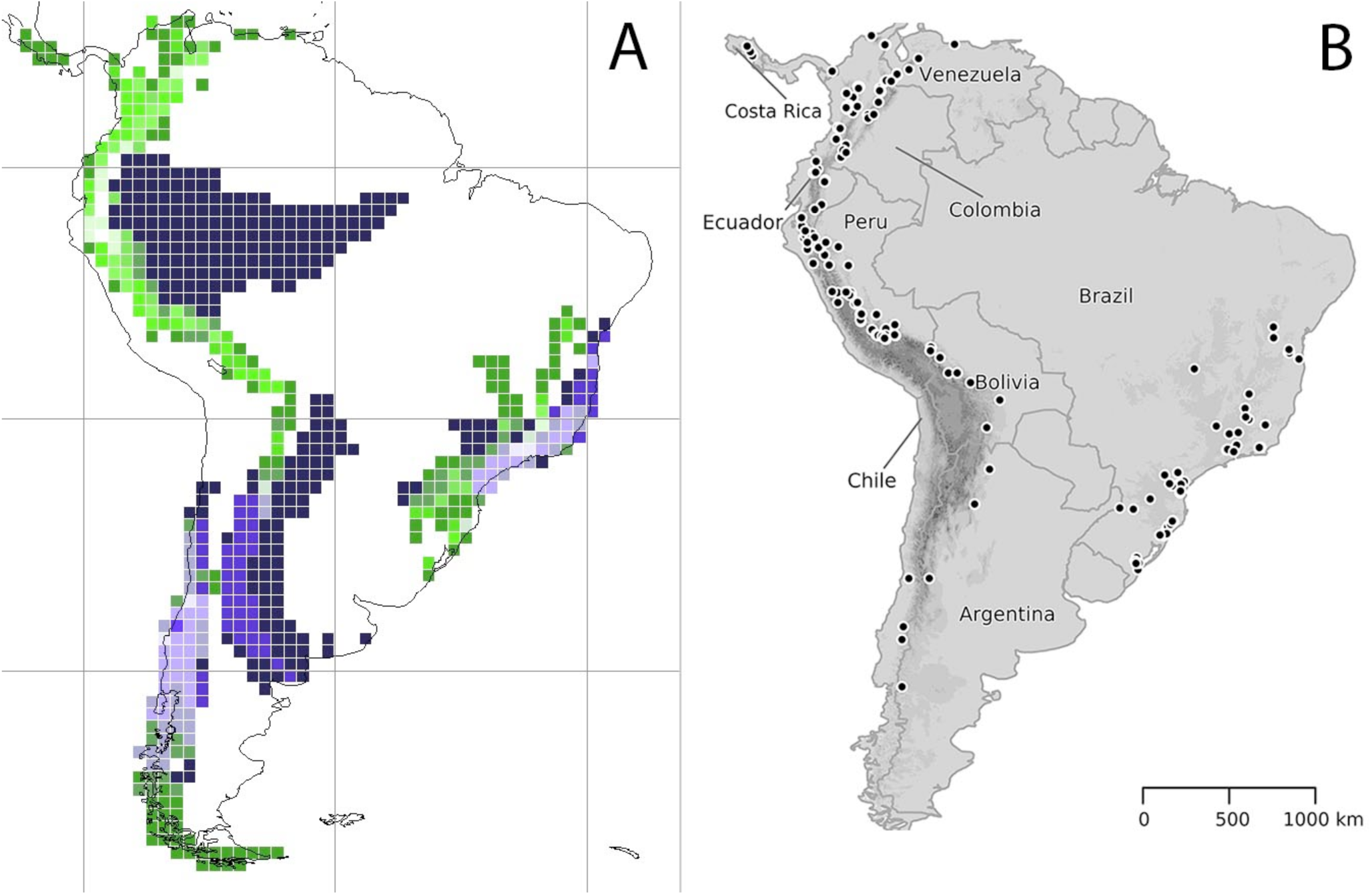
(a.) Geographic pattern of species richness in the family Rhinocryptidae in a grid corresponding to 1×1 degrees; brightness reflects species richness, from dark blue areas with only one species to a white cell in northern Ecuador with nine species. Species in genera different from *Scytalopus* (corresponding roughly to the first root-path quartile in the family phylogeny, or the 25% of species with few or no near relatives) are represented by purplish blue hues, and species of *Scytalopus* are represented by green hues (producing greyish hues where the two categories overlap). (b.) Localities where specimens of *Scytalopus* were sampled for molecular phylogenetic analyses in this study; maps showing geographic sampling per species are in the Supplementary Material.

Aside from its biogeographic relevance as a model to study diversification in the montane Neotropics, *Scytalopus* is of particular interest from a systematics perspective because it is among the most challenging avian genera for understanding species limits. Because of their retiring habits, *Scytalopus* tapaculos are difficult to observe and collect (Chapman 1915), and species are remarkably similar morphologically. Many taxa cannot be distinguished from each other even in the hand, which explains why traditional museum-based classification lumped populations inhabiting equivalent ecological zones in different mountains into broadly defined biological species (Zimmer 1939). In the 1990s, a shift to a modern taxonomic approach integrating information from specimens, vocalizations, and geographical and ecological distributions resulted in considerable advances in our understanding of species limits in the group (Whitney 1994; Fjeldså and Krabbe 1990; Krabbe and Schulenberg 1997; Vielliard 1990). These efforts revealed that previously overlooked species replace each other sharply in different elevational zones and on different mountain slopes, and set the stage for additional assessments of the taxonomic status of populations as well as descriptions of numerous new taxa (Avendaño et al. 2015; Avendaño and Donegan 2015; Bornschein et al. 2007; Bornschein et al. 1998; Coopmans et al. 2001; Cuervo et al. 2005; Donegan et al. 2013; Donegan and Avendaño-C. 2008; Hosner et al. 2013; Krabbe and Cadena 2010; Krabbe et al. 2005; Maurício 2005; Maurício et al. 2014; Raposo et al. 2006; Stiles et al. 2017; Whitney et al. 2010). Although more work is required to arrive at a classification of *Scytalopus* reflecting its true diversity, progress has been substantial: whereas Zimmer (1939) recognized 10 species in the genus –some of which are not considered valid or part of the genus any more–, current taxonomic treatments recognize 44 (Remsen et al. 2018; Krabbe and Schulenberg 2003; Gill and Donsker 2019).

Given the apparent uniformity in morphology and coloration across *Scytalopus* and the incomplete knowledge of their diversity, molecular data can be useful to reveal genetically divergent lineages and potentially cryptic species. Accordingly, crucial to recent advances in the systematics of the group has been the advent of DNA sequences used to establish the genetic distinctiveness and phylogenetic affinities of taxa (e.g., Mata et al. 2009; Pulido-Santacruz et al. 2016; Avendaño et al. 2015; Krabbe and Cadena 2010; Stiles et al. 2017). Coupled with anatomical data, molecular characters further served to reveal a striking discordance between external appearance and phylogeny, which led to the separation of *Scytalopus* as traditionally defined in two separate genera, one of which (true *Scytalopus*) is in a clade with *Myornis* and *Eugralla*, whereas the other (*Eleoscytalopus*) is more closely allied to *Merulaxis* (Maurício et al. 2008). Within the former clade, it remains unclear whether *Scytalopus* as recently redefined is monophyletic with respect to *Myornis* and *Eugralla* because few *Scytalopus* species have been considered in analyses (Maurício et al. 2008; Ohlson et al. 2013; Ericson et al. 2010; Maurício et al. 2012). In fact, the only phylogenetic study available for *Scytalopus* including more than a handful of species from the Andes was based only on 285 base pairs of mitochondrial DNA sequence for taxa occurring in Ecuador and Peru (Arctander and Fjeldså 1994), and some of its results were compromised because nuclear pseudogenes were considered in analyses (Arctander 1995). Most other molecular studies of *Scytalopus* have concentrated on few taxa relevant for taxonomic assessments (Cuervo et al. 2005; Krabbe and Cadena 2010; Krabbe et al. 2005; Avendaño et al. 2015; Stiles et al. 2017), and notably on the phylogeography and diversification of Brazilian taxa (Mata et al. 2009; Pulido-Santacruz et al. 2016). Data for Brazilian and Andean species have not been integrated in comprehensive analyses of phylogeny and diversification.

We used DNA sequences from nuclear and mitochondrial loci to construct a robust phylogeny of *Scytalopus* including all described species, nearly all named forms, several unnamed taxa, and multiple individuals representing geographic variation within species. We employed this data set to assess the monophyly of *Scytalopus* and to examine the relationships among species and species-groups in a geographic context, asking whether the tropical Andes were recruited upslope from adjacent lowlands or from the cool austral parts of South America. We also used relaxed molecular clock methods and estimates of diversification rates of clades through time to examine the timing and mode of diversification in the group aiming to determine whether Andean species diversity (1) was built up gradually in association with geological uplift, (2) was punctuated by events such as colonization of new regions, (3) or followed cycles of fragmentation and reconnection of populations in response to Pleistocene glacial cyles. We also examined the accumulation of lineages over time as a first approximation to assessing whether diversification may have been constrained by filling of ecological space. Finally, because species limits in *Scytalopus* remain unclear and taxonomy likely underestimates its true diversity, we employed our molecular data to highlight divergent lineages which may represent distinct species.

## Methods

### Taxon and Gene Sampling

Current taxonomy recognizes 44 species of *Scytalopus* (Remsen et al. 2018; Gill and Donsker 2019), all of which were included in our analyses (Appendix 1). We also sampled museum specimens of all but one named forms (*S. argentifrons chiriquensis* from western Panama), attempting to analyze specimens from near the type locality of taxa whenever possible. We sought to represent intraspecific variation by sampling multiple specimens per species. This approach resulted in a sampling scheme including up to >20 individuals per named species as well as representatives of undescribed species including populations from Pasco, Junín, Ayacucho, and Lambayeque-Cajamarca, Peru (Fjeldså and Krabbe 1990; Krabbe and Schulenberg 2003; Hosner et al. 2015). To test the monophyly of *Scytalopus* based on broader taxon sampling than in previous studies, we also included the monotypic *Myornis senilis* and *Eugralla paradoxa* as part of our ingroup. Based on relationships documented by earlier work (Irestedt et al. 2002; Maurício et al. 2008; Ohlson et al. 2013; Ericson et al. 2010), we used taxa in the genera *Rhinocrypta, Merulaxis, Eleoscytalopus, Acropternis, Scelorchilus,* and *Pteroptochos* as outgroups.

We generated two data sets to examine relationships among *Scytalopus* species and close relatives. First, we sequenced a c. 1000 bp fragment of the second subunit of the NADH dehydrogenase mitochondrial gene (ND2) for most available samples, resulting in extensive geographic coverage of the montane Neotropics (Figure 1). The majority of the ND2 sequences of Andean specimens we employed are new, but we reported a few of them previously in focused taxonomic studies (references and GenBank accession numbers in Appendix 1). All sequences of Brazilian specimens were obtained from the literature (Mata et al. 2009; Pulido-Santacruz et al. 2016; Appendix 1). In total, we analyzed ND2 data for a total of 310 individuals: 304 *Scytalopus,* 1 *Myornis,* 1 *Eugralla*, and 4 outgroups.

Guided by taxonomy, by results of preliminary analyses of ND2 sequences, and by an ongoing study reconstructing a species-level phylogeny of the suborder Tyranni (M. Harvey et al., in prep), we selected 63 specimens representing 42 named species and 11 subspecies of *Scytalopus* (missing species were *S. alvarezlopezi* and *S. simonsi*), as well as four individuals in the genera *Rhinocrypta, Merulaxis, Myornis*, and *Eugralla* for multilocus assessments of relationships (Appendix 2). This strategy involved a sequence capture approach targeting ultraconserved elements (UCEs; Faircloth et al. 2012) and conserved exons across the genome (Zucker et al. 2016). After quality control, trimming, assembly, and filtering this dataset comprised a total of 1,246,332 bp distributed in 1,833 UCEs (1,201,528 bp) and 80 exons (44,804 bp).

We extracted total DNA from frozen tissues using standard methods and kits (Qiagen, Valencia, CA, and Invitrogen, Carlsbad, CA). DNA extraction from toepads was conducted in facilities dedicated to work with ancient DNA to avoid contamination. Protocols for DNA extraction from toepads followed recommendations from comercial kits, but they included longer digestion times with the addition of dithiothreitol (DTT). ND2 was amplified and sequenced using standard methods (Cadena et al. 2007). We quantified DNA concentration using a QuBit fluorometer (ThermoFisher, Waltham, MA) and sent DNA extracts to Rapid Genomics (Gainesville, FL) for library preparation and sequence capture following the general protocol described by Faircloth et al. (2012).

### Phylogenetic analyses

We used the nuclear UCE and exon data set to provide an overall assessment of phylogenetic relationship among species of *Scytalopus* and near relatives. We extracted sequences for the 63 rhinocryptid individuals included in analyses from sequence alignments built using the Phyluce pipeline (Faircloth 2016) as part of a species-level phylogenomic analysis of the suborder Tyranni (M. Harvey et al. in prep). We realigned sequences using MAFFT v.7.310 (Katoh and Sandley 2013) and estimated phylogenetic hypotheses following two strategies. First, we conducted a concatenated maximum-likelihood analysis partitioned by locus type (i.e., exons and UCEs) using the GTRCAT substitution model in RAxML v8.2.10 (Stamatakis 2006). Second, after removing two samples with the highest proportion of missing data across loci, we conducted species-tree analyses using ASTRAL II v.4.10.12 (Sayyari and Mirarab 2016). We estimated maximum-likelihood gene trees for each locus using the GTRGAMMA model of nucleotide subsitution using RAxML. We performed 100 bootstrap replicates per locus, which also allowed us to run ASTRAL performing 100 bootstrap replicates.

For the mtDNA data set, we conducted maximum-likelihood analyses using RAxML v. 8 (Stamatakis 2014), and also employed Bayesian approaches implemented in MrBayes v3.3.2 (Ronquist et al. 2012) and Beast 2.4.8 (Bouckaert et al. 2014) to estimate tree topologies and divergence times. We conducted the maximum-likelihood analyses using RaxML under a GTR+Γ model of nucleotide substitution with 1,000 bootstrap replicates, specifying separate partitions by codon. The Bayesian analysis in MrBayes was conducted implementing a partition scheme by codon position as suggested by MrModeltest 2.3 (Nylander 2004), and consisted of four MCMC chains of 20 million generations, sampling every 1,000 generations, and discarding the first 50% as burn-in. For the Bayesian analysis conducted using Beast we ran 100 million generations, sampling trees and parameters every 5000 generations. A relaxed uncorrelated clock (lognormal distribution, mean = 0.0125, SD = 0.1; Smith and Klicka 2010) and a birth-death speciation tree prior (Ritchie et al. 2017) were applied. We also conducted the Beast analysis with an alignment of 90 sequences (83 *Scytalopus*) running a chain of 100 million generations and sampling trees and parameters every 5000 generations. These 83 sequences were chosen based on our best (albeit subjective) judgement of the existence of distinct taxa within *Scytalopus* based on current taxonomy, our knowledge of geographic variation in vocalizations, and genetic structure. We applied a relaxed uncorrelated clock and a Yule speciation tree prior. We confirmed likelihood stationarity and adequate effective sample sizes above 500 for all estimated parameters using Tracer v1.6.0 (http://tree.bio.ed.ac.uk/software/tracer). The parameter values of the samples from the posterior distribution on the maximum clade credibility tree were summarized after discarding the first 50% as burn-in using TreeAnnotator v1.10.1 (Drummond et al. 2012). We ran all analyses through the CIPRES Science Gateway V. 3.3 (Miller et al. 2010).

### Diversification through time

Based on our time-calibrated mtDNA phylogeny, we explored temporal patterns of diversification in *Scytalopus*. First, we examined the accumulation of lineages along the history of the group by constructing lineage-through-time (LTT) plots, which show the (log) cumulative number of lineages as a function of time. We constructed plots based on the maximum clade credibility tree obtained in Beast, and also across a sample of 100 trees in the posterior distribution to consider uncertainty in phylogenetic inference. Second, to determine whether speciation in *Scytalopus* occurred predominantly during the relatively recent past of the group as expected if speciation was triggered by Pleistocene glacial cycles, or if it concentrated early on in the history of the group as predicted by the hypothesis that diversification was driven by ecological opportunity, we calculated the gamma statistic (Pybus and Harvey 2000). When gamma takes high positive values, one can infer an increasing rate of diversification towards the present, whereas high negative values indicate a slowdown of the diversification rate.

We also used the program BAMM (Bayesian Analysis of Macroevolutionary Mixtures v 2.5.0; Rabosky et al. 2013) to examine shifts in rates of diversification across clades and over time. We ran the analysis 10 times, each for 1,000,000 generations, and sampled one of every 1,000 trees generated during each run. The evolutionary rate priors were calculated using the setBAMMpriors function in the R package BAMMtools (v2.1.6; Rabosky et al. 2014). Also using BAMMtools, we confirmed that all ESS values were above 500, signaling convergence of parameter estimates.

Given uncertainty about true species diversity in *Sctytalopus,* we explored the sensitivity of our inferences based on the above diversification analyses to alternative taxonomic schemes. We separately built LTT plots, estimated the gamma statistic, and conducted BAMM analyses for two different data sets obtained by excluding specimens from the Beast tree consisting of 90 sequences described above. First, in a 48-tip data set, we trimmed specimens from the tree so that terminals for analyses were the 44 currently accepted species in the genus plus four unnamed species vocally distinct from others to be described in forthcoming publications (N. Krabbe et al. and J. Schmitt et al., in preparation). Second, we analysed an 83-tip data set consisting of representatives of the 48 taxa above, plus lineages we recognize as distinct based on genetic differentiation and our knowledge of geographic variation in plumage and vocalizations.

## Results

### Phylogeny and biogeography

Analyses using alternative data sets (1,833 UCE loci and 80 nuclear exons for 63 terminals, ND2 gene for 311 terminals) and methods of phylogenetic inference (maximum-likelihood and Bayesian inference of gene trees, species trees) produced congruent results (Figures 2-4). Across all analyses, we found strong support for the monophyly of *Scytalopus,* with the austral *Eugralla* as its sister group, and the tropical Andean *Myornis* sister to the *Scytalopus-Eugralla* clade. Within *Scytalopus,* the deepest split was between two strongly supported clades. The first clade included all of the Brazilian species, which was sister to a clade comprising almost exclusively species from southern South America (oligothermic areas of southern Peru, Bolivia, Argentina, and Chile; hereafter Southern Andes clade). The other main clade included nearly all taxa occurring in the northern sector of the distribution of the genus, from Costa Rica to Central Peru (hereafter Tropical Andes clade). The clearest exception to these general patterns was that members of the *S. canus* complex (i.e. *S. canus, S. opacus*), from paramos of the northern Andes, were embedded in the Southern Andes Clade and not in the Tropical Andes clade where all other northern taxa belonged. In addition, *S. parvirostris* and *S. bolivianus*, two species in the Tropical Andes clade, reach south to the zone of winter chills in the Bolivian Yungas.

**Figure 2.**
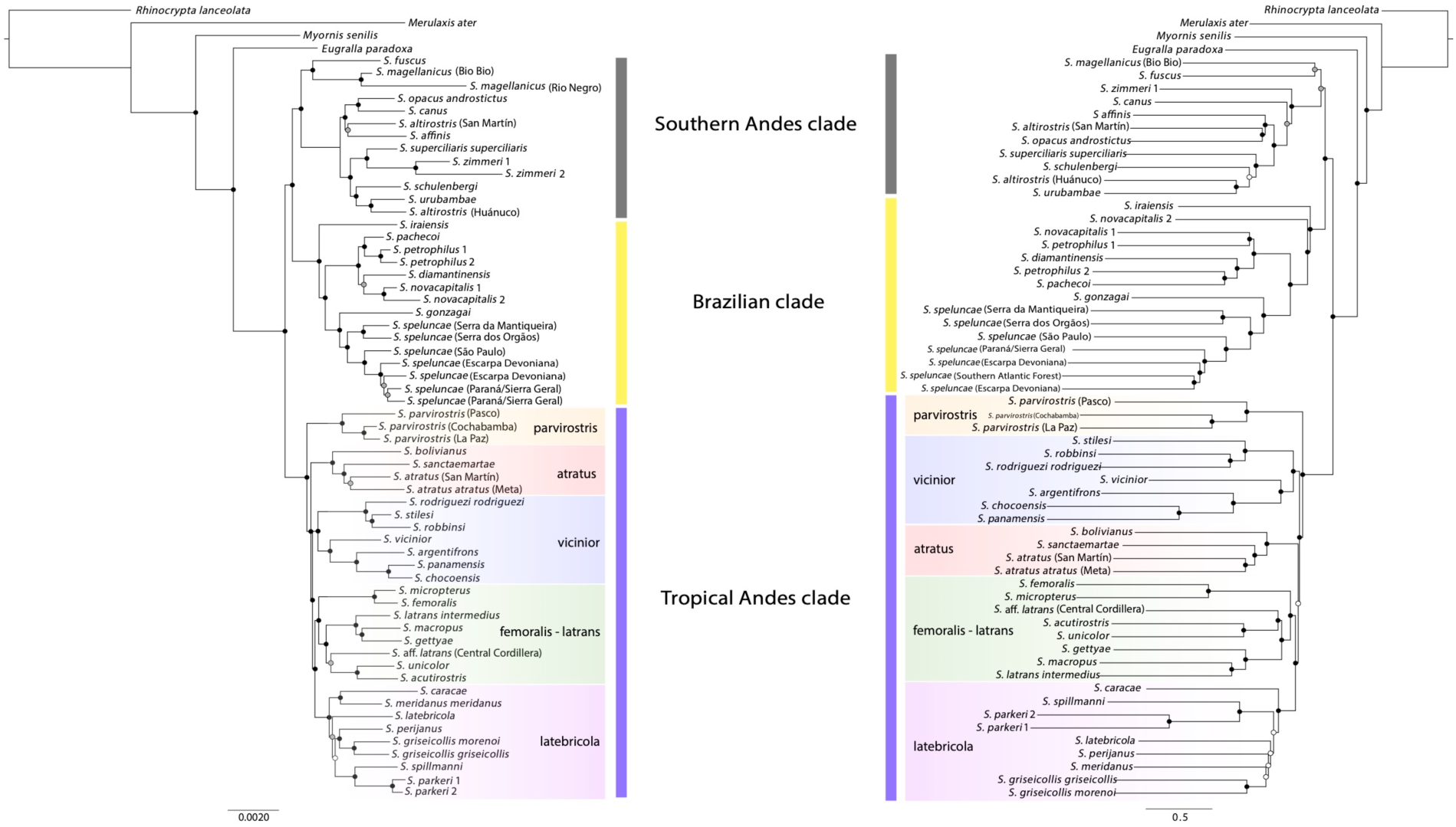
Phylogenetic relationships of *Scytalopus* tapaculos and allies inferred based on concatenated maximum-likelihood (left) and species-tree analyses (right) of 1,833 UCEs and 80 exons in the nuclear genome. The results are congruent in documenting the existence of three strongly supported groups associated with geographic regions: (1) a Southern Andes clade, (2) a Brazilian clade, and (3) a Tropical Andes clade. Traditionally recognized species complexes in the Tropical Andes were recovered with high support in both analyses. The color of circles on nodes indicates bootstrap support values: <70% (white), 70 – 90% (gray), and >90% (black).

The deepest split within the Southern Andes clade separated two sister species with the southernmost geographic ranges in the genus (*S. magellanicus* and *S. fuscus*) from a clade formed by all other taxa (Figures 2 and 3). The Brazilian clade consisted of two main monophyletic groups: one group included a clade formed by various lineages of *S. speluncae*, which was sister to *S. gonzagai*, whereas the other group included *S. iraiensis* and a clade formed by *S. novacapitalis, S. pachecoi, S. diamantinensis*, and *S. petrophilus*. Within the Tropical Andes clade, all analyses recovered five distinct and well supported clades, which we refer to as the *parvirostris, atratus, vicinior, femoralis,* and *latebricola* groups. Relationships among those groups were not entirely clear because branches were short and topologies differed slightly between analyses employing different methods (i.e. concatenated ML vs. species-tree in nuclear data) and data sets (i.e. nuclear vs. mitochondrial DNA). The *parvirostris* group, which as a whole has the southernmost distribution in the clade (the species *S. bolivianus* in the *atratus* group ranges farther south), was sister to a clade formed by all other groups from the Tropical Andes in trees based on nuclear loci (Figure 2) but this relationship was not supported in the ND2 trees (Figures 3 and 4).

**Fig. 3.**
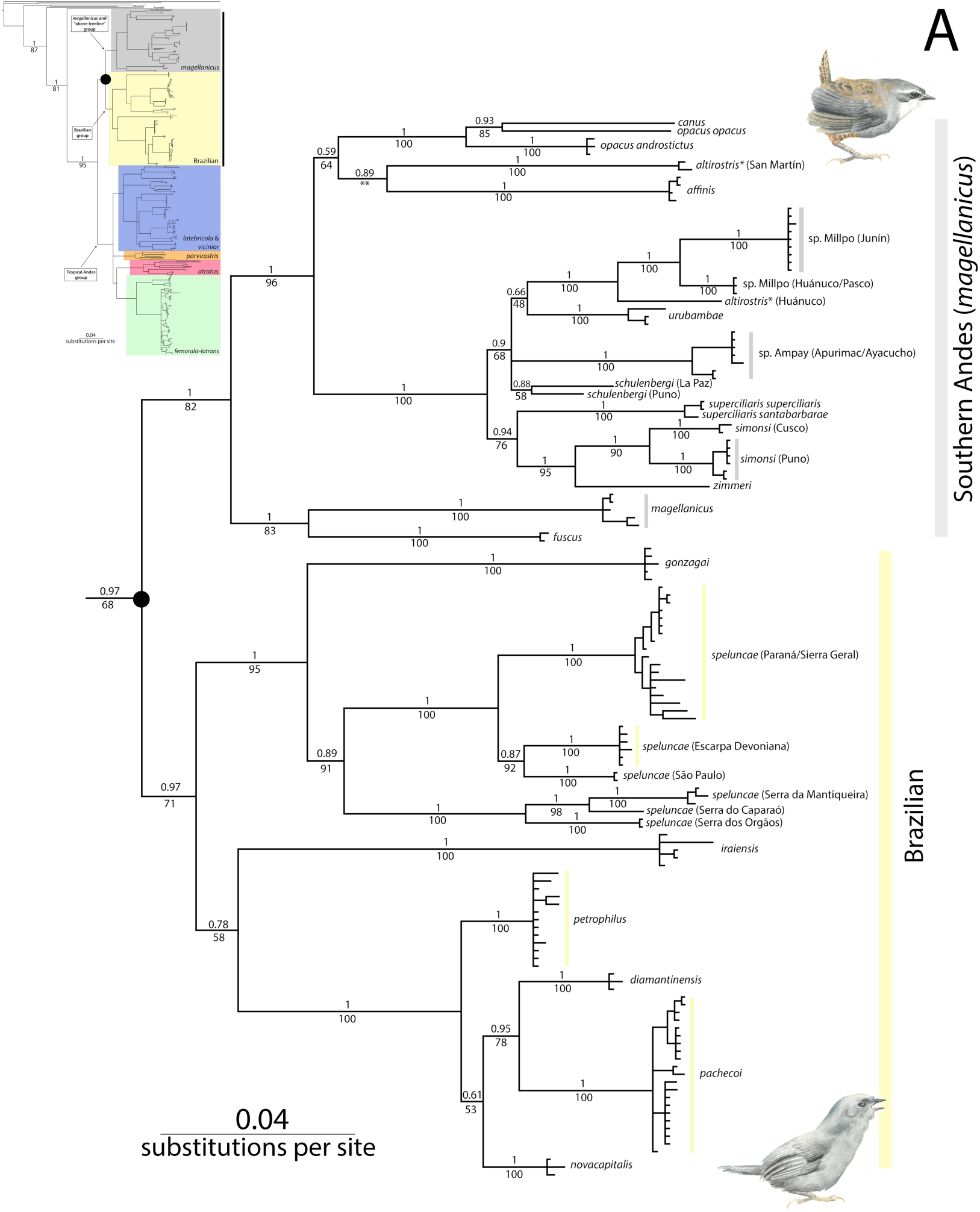
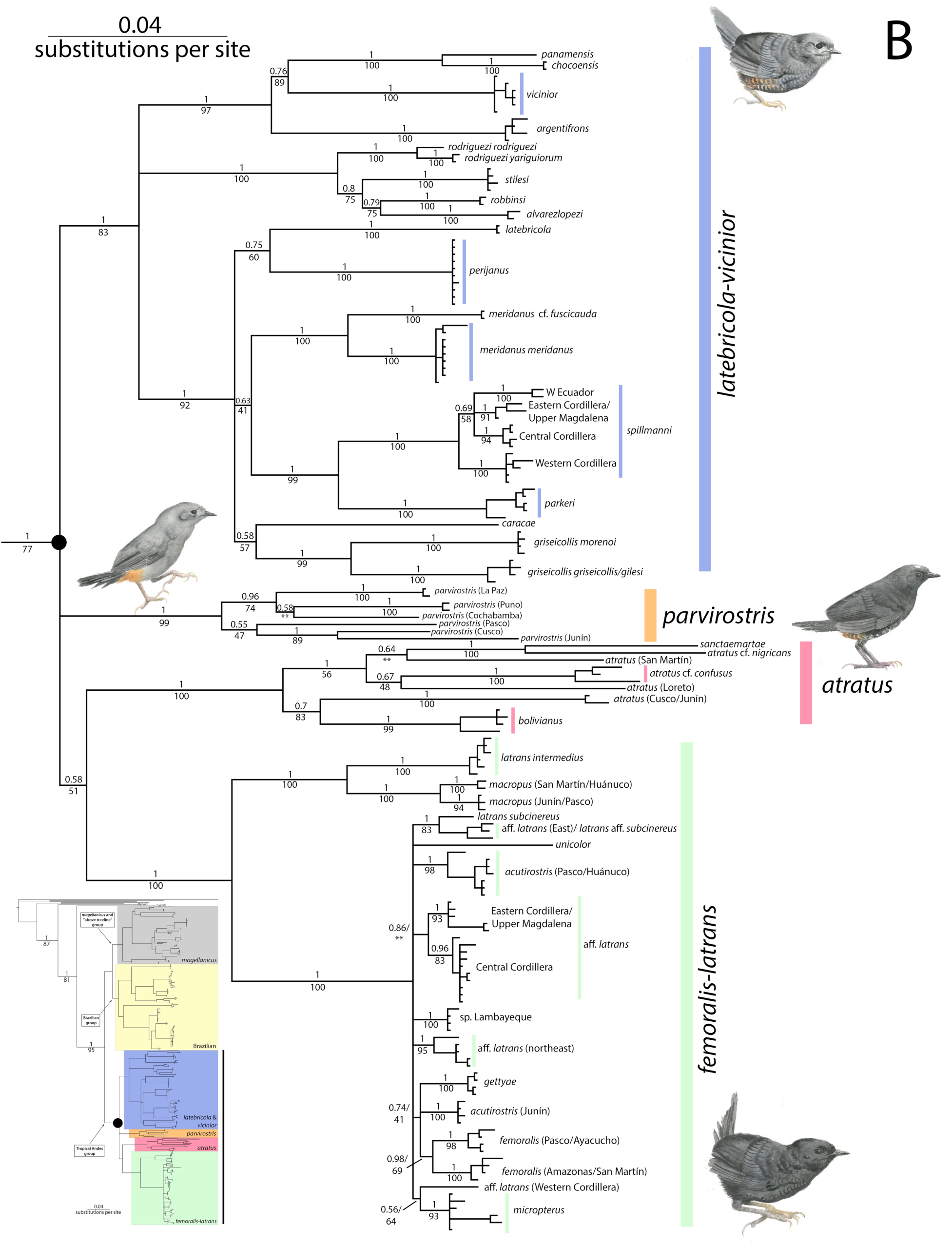
Bayesian tree showing phylogenetic relationships of *Scytalopus* tapaculos and allies inferred using MrBayes based on sequences of the ND2 mitochondrial gene. (A) focuses on a clade formed by sister groups from mostly the Southern Andes (*magellanicus* and “above treeline” clade) and eastern Brazil, whereas (B), shown in the following page, focuses on relationships within the Tropical Andes clade, which is sister to the clade in (A). Posterior probabilities are shown above branches and maximum-likelihood bootstrap values obtained using RaxML below branches. Double asterisks are shown for clades not recovered in the maximum-likelihood analysis. The small tree in each panel shows relationships of major clades, with circles and vertical bars identifying clades shown in detail in each case. Illustrations depict a representative species in each of the major clades: (A.) *Scytalopus zimmeri* for the *magellanicus* group and *S. speluncae* for the Brazilian group, (B.) *S. vicinior* for the *latebricola* + *vicinior* group, *S. parvirostri*s, *S. atratus*, and *S. latrans* for the *femoralis-latrans* group. Watercolors by Jon Fjeldså.

**Figure 4.**
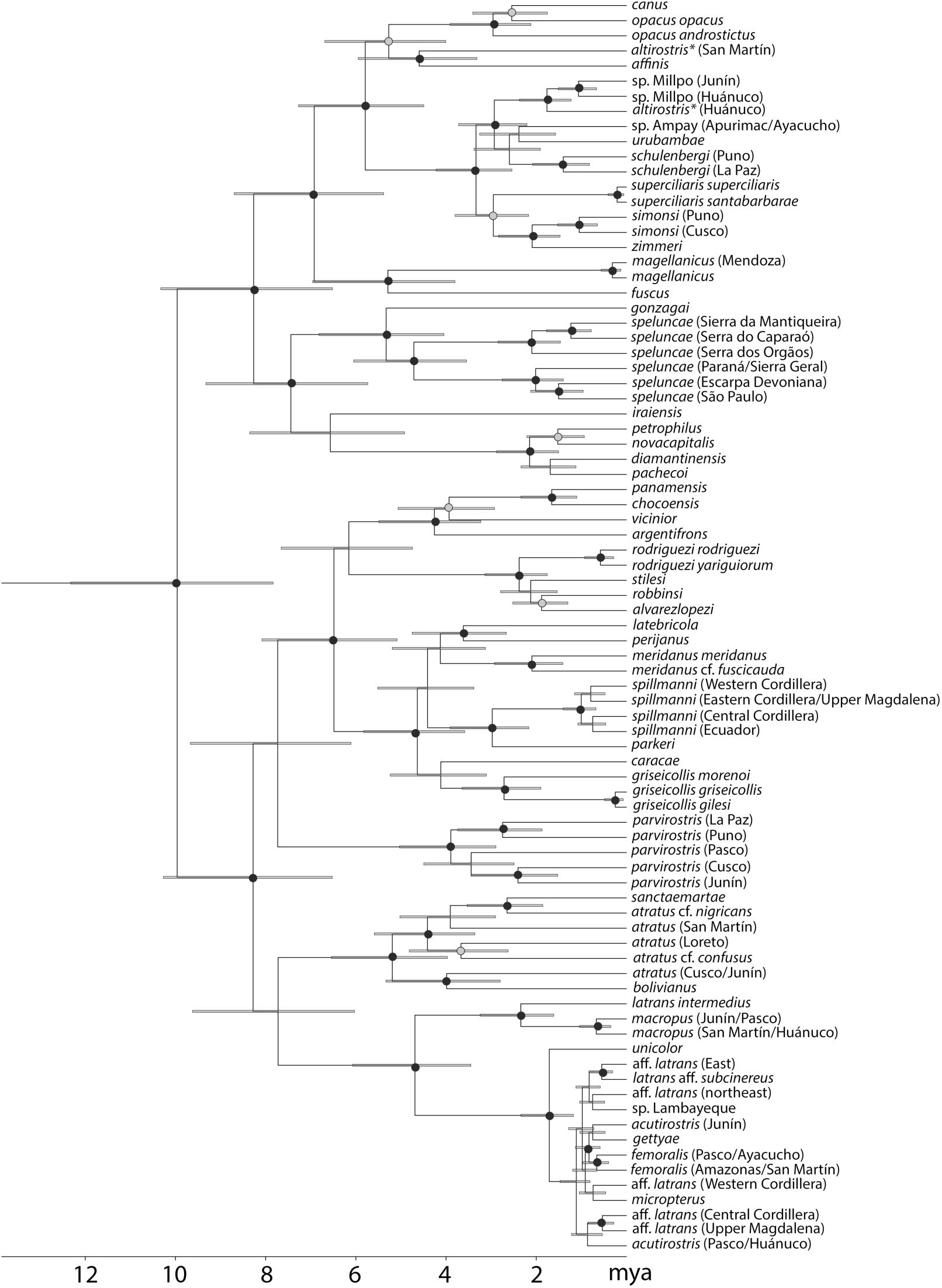
ND2 gene tree estimated with Beast showing divergence times in million years before present within *Scytalopus*. Gray bars correspond to the 95% highest posterior density (HPD) intervals for the ages of nodes. Dots on nodes indicate posterior probabilities when ≥ 0.70: grey 0.70-0.90, black >0.90.

The majority of species for which we sampled multiple individuals for mtDNA analyses were reciprocally monophyletic (Figure 3). One exception was *S. atratus*, a widespread species showing substantial geographic structure, which was paraphyletic with respect to *S. sanctaemartae* and *S. bolivianus*. A second exception was *S. latrans*: subspecies *S. l. intermedius* was closely allied to *S. macropus*, whereas subspecies *S. l. subcinereus* and nominate *S. l. latrans* belonged to a clade also including *S. unicolor, S. femoralis, S. gettyae, S. micropterus, S. acutirostris*, and an undescribed species from Lambayeque and Cajamarca, Peru. Relationships among these taxa, among populations of nominate *S. l. latrans*, and among different populations of other species (*S. acutirostris, S. femoralis*) were unresolved, likely a consequence of recent and rapid divergence. Third, birds sampled in Peru referred to *S. altirostris* did not form a clade and were, in fact, distantly related to each other based both on the nuclear (Figure 2) and mitochondrial (Figures 3 and 4) data. One individual from San Martín appeared most closely allied to *S. opacus, S. canus*, and *S. affinis*, whereas an individual from Huánuco north of the Río Huallaga formed a clade with *S. urubambae* and undescribed taxa from Junín and Huánuco south of the Huallaga. Finally, Peruvian specimens identified as *S. acutirostris* formed two distinct clades, one occurring in Pasco and Huánuco, the other in Junín; although the phylogenetic position of such clades was not strongly supported, they were not recovered as sister in any analysis of mitochondrial data (Figures 3 and 4; nuclear data was obtained from a single individual).

Genetic structure within named species largely reflected geography (Figure 3). For example, within *S. spillmanni*, a species in which no geographic variation in plumage has been recognized at the subspecies level, birds from each of the three cordilleras of the Colombian Andes formed distinct clades; birds from Ecuador either formed a separate group or belonged to the clade from the Western Cordillera of Colombia. Surprisingly, we uncovered deep genetic divergence among populations of the monotypic *S. parvirostris*. Except for a genetic break observed between Cusco and Puno in southern Peru (i.e. a region lacking overt habitat discontinuities; Cadena and Cuervo 2010), genetic structure within *S. parvirostris* seemed to be associated with dry enclaves bisecting the cloud forest belt along the eastern slope of the Peruvian and Bolivian Andes, including the Apurímac, Urubamba, and Cotacajes valleys. Likewise, a deep split in *S. griseicollis* existed between populations from the northern and southern parts of its range in the Colombian Cordillera Oriental; this split coincided with divergence between subspecies *S. g. morenoi* and a clade including nominate *S. g. griseicollis* and *S. g. gilesi* (Avendaño and Donegan 2015). Divergence within the Venezuelan endemic *S. meridanus* also reflected geography, with distinct clades occurring in different regions within the Cordillera de Mérida corresponding to subspecies *S. m. meridanus* and *S. m.* cf. *fuscicauda*. Finally, marked population structure associated with geography was also observed within *S. speluncae* in Brazil as previously documented (Pulido-Santacruz et al. 2016).

### Diversification through time

Our time-calibrated tree (Figure 4) indicated that *Scytalopus* diverged from its sister group in the Mid Miocene and began diversifying into the main clades formed by extant species some 10 m.a. (crown age 7.8-12.3 m.a. highest posterior density, HPD). Divergence between the Southern Andes and Brazilian clades occurred ca. 8 m.a. (6.5-10.3 m.a. HPD), roughly coinciding with early divergence events within the Tropical Andes clade. Divergence between the only Central American member of the genus (*S. argentifrons* from Costa Rica and western Panama) and its sister group (a clade formed by *S. chocoensis, S. panamensis*, and *S. vicinior*) was estimated to have occurred ca. 4 m.a. (3.2-5.5 m.a. HPD).

Lineage-through-time plots (Figure 5A) revealed roughly constant accumulation of diversity through the Late Miocene and Pliocene within *Scytalopus*, with an overall apparent downturn in lineage accumulation in the Pleistocene reflected in negative values for the gamma statistic in the complete phylogeny and within its main clades (Figure 5B). Based on the 83-tip data set, diversification rates estimated using BAMM appeared to have steadily declined since *Scytalopus* originated, with a notable exception being a significant upward shift in diversification rate within the *femoralis-latrans* group, which diversified into numerous taxa in the Pleistocene beginning ca. 1.5. m.a. (Figure 5C, 5D). The two rate-shift configurations with highest posterior probabilities involved a single shift leading to increased diversification rates within the *femoralis-latrans* group. In one case, the clade with increased rates was defined by the most recent common ancestor of *S. unicolor* and *S. femoralis* (posterior probability 0.54, Figure 5C), whereas in the other (posterior probability 0.29) *S. unicolor* was not part of the clade with increased rates. A configuration with no shifts in rates had a lower posterior probability (0.16). In contrast, when we conducted analyes based on the 48-tip data set, we found no support for shifts in diversification rates within *Scytalopus*; the configuration with the highest posterior probability (0.77) did not include shifts in diversification rates, and configurations having rate shifts in clades matching those identified in the analysis with 83 tips had much lower support (0.17 and 0.04 posterior probabilities). We observed no significant changes in diversification rates potentially linked with the colonization of new regions.

**Figure 5.**
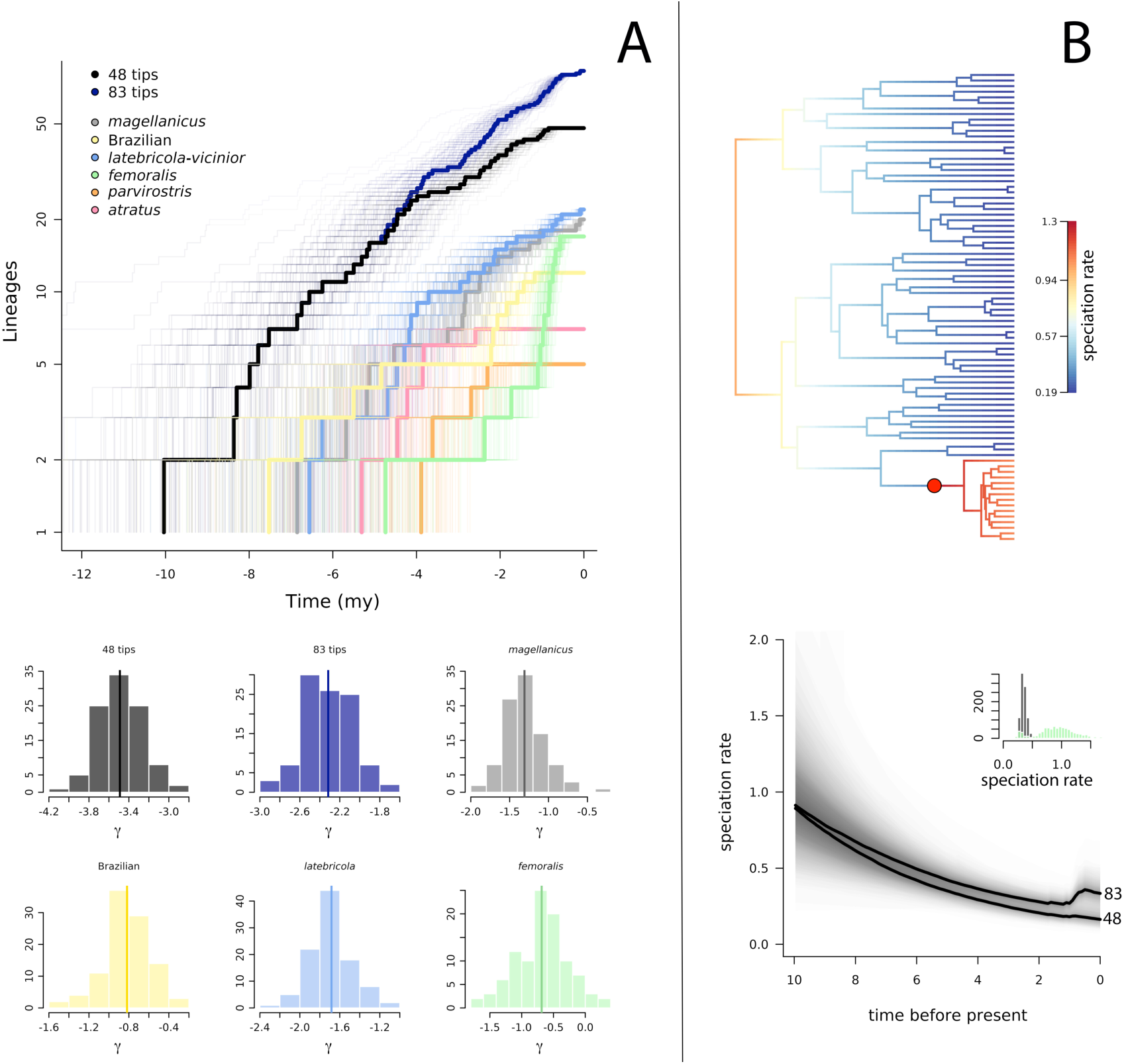
Analyses of diversification dynamics in *Scytalopus* tapaculos reveal declining rates in the accumulation of lineages over time across the group, except in one of the clades from the Tropical Andes. (A, top) Lineage-through-time plots show a downturn in lineage accumulation for the whole genus regardless of whether one analyzes data only for described and soon-to-be-described species (48 tips), or employs a more liberal approach treating lineages differing vocally or genetically as distinct (83 tips). Downturns in lineage accumulation are also observed within major clades. Thick lines are plots based on the maximum clade credibility tree obtained using Beast and thin lines are from 100 trees in the posterior distribution in each case. (A, bottom) Estimates of the gamma statistic are negative across the phylogeny and within all clades confirming declining diversification rates over time. Histograms show the frequency distribution of gamma across trees in the posterior and vertical lines indicate median estimates of gamma based on the 83-tip data set. (B, top) Instantaneous rates of speciation estimated using BAMM mapped onto the Beast tree constructed using the 83-tip data set; speciation rates declined consistently across the phylogeny, with a clade in the *femoralis* group (indicated with red circle) being the only clade exhibiting a significantly upward shift in speciation rate. (B, bottom) Declining speciation rate across the phylogeny through time as estimated with both data sets, with an inferred increase in the past ca. 1.5 million years detected when employing the 83-tip data set reflecting the increase in rate within the *femoralis* clade. The thick lines depict the mean across samples in the posteriors estimated by BAMM; the histogram in the inset shows the posterior distribution of speciation rate across the tree (black) and in the *femoralis* clade (green) based on the 83-tip data set.

## Discussion

### Phylogeny and biogeography of Scytalopus *tapaculos*

The origin of the montane fauna of northern South America hypothetically followed the uplift of the Northern Andes, which created new habitats (i.e. stunted cloud forest and subsequently subalpine páramo) colonized by lineages originating in temperate latitudes tracking their favored environmental conditions (Vuilleumier 1986; Hoorn et al. 2013; Kattan et al. 2016; Chapman 1917). Most of the deep branches in the Rhinocryptidae phylogeny (Ericson et al. 2010), which diverged in the Oligocene and early Miocene (Ohlson et al. 2013), are found in southern South America up to the Brazilian Atlantic Forest, with three independent colonizations extending further north (*Lioscelis thoracicus* in the Amazon, and *Acropternis orthonyx* and *Myornis senilis* in the northern Andes). In agreement with anatomical data (Maurício et al. 2012), our results clearly support the monophyly of *Scytalopus* with respect to *Eugralla* and *Myornis.* Our analyses further indicate that *Scytalopus* radiated rapidly from the Late Miocene, apparently with a northwards progression and with most speciation taking place well before the onset of large-amplitude climatic oscillations in the Late Pleistocene (Hooghiemstra and Van der Hammen 2004).

The sister-group relationship between a Southern Andes clade and a Brazilian clade we uncovered in *Scytalopus* conforms with a pattern seen in other avian groups with representatives in the Andes and eastern Brazil (Silva 1995). Sick (1985) and Vielliard (1990) suggested a single colonization by tapaculos from the Andes to eastern Brazil but two of the *Scytalopus* species they recognized were found to be related to *Merulaxis* and are now placed in *Eleoscytalopus* (Maurício et al. 2008). Additional discussion has focused on the route through which tapaculos may have dispersed between the Andes and the Brazilian highlands (Sick 1985; Vielliard 1990; Maurício 2005; Willis 1992). Our estimates of divergence times indicate that *Scytalopus* sensu stricto was established in southeastern Brazil by the Late Miocene, after the geological collision between the Brazilian Shield and the ‘Chapare Buttress’ of the rising Andes gave rise to a continuum of rugged and forest covered landscapes. During the final uplift of the Altiplano, geological subsidence in the Beni Plains and Chaco gave rise to hydrologically unstable low plains (Hanagarth 1993), which may have been unsuitable for tapaculos, presumably leading to extinctions and subsequent isolation of the Brazilian populations. Because other researchers are working on biogeography and diversification of Brazilian tapaculos (G. Maurício et al. in litt.), we do not further discuss patterns in that region. However, we do note that phylogenetic and phylogeographic studies have estimated considerably more recent divergence times between Andean and Brazilian lineages than those we estimated for *Scytalopus* (García-Moreno et al. 1999; Trujillo-Arias et al. 2017; Trujillo-Arias et al. 2018; Cabanne et al. 2019), suggesting that biogeographic connections through the Chaco or Cerrado for montane taxa likely existed at various moments in the past (see also Prates et al. 2017).

The species of *Scytalopus* reaching the highest elevations in the Northern Andes are members of the Southern Andes clade (*S. canus, S. o. opacus, S. o. androstictus*), which are restricted to microphyllous vegetation at treeline in Colombia, Ecuador, and northwestern Peru (Krabbe and Cadena 2010). The phylogenetic position of the *S. canus* complex within a clade formed by species occurring in the southernmost areas of South America occupied by the genus (from the southern tip of the continent through Bolivia and Peru) and its distant relationship to other Northern Andean taxa (i.e. the Tropical Andes clade), is thus consistent with the hypothesis that various organisms colonized the high elevations of the Northern Andes from temperate regions (Cadena 2007; Vuilleumier 1986; Hoorn et al. 2013); a similar pattern has also been observed in plants from the Andes (Hughes et al. 2013) and African highlands (Gehrke and Linder 2009). Niche conservatism may partly account for the high species richness of the tropics because lineages which evolved when global climate was generally warm are unable to colonize temperate latitudes (or highlands) because they cannot adapt to cool temperatures (Wiens and Donoghue 2004; Fine 2015). Our results add a twist to this niche conservatism hypothesis in suggesting that ecological conservativeness of temperate-zone lineages which manage to colonize cool tropical environments and diversify within them (i.e. the *S. canus* group in the Northern Andes) may also contribute to the high diversity of tropical areas (Moriniére et al. 2016; see also Bacon et al. 2018). Although clades of high-latitude origin may more easily tolerate tropical conditions (Smith et al. 2012; Araújo et al. 2013; Khaliq et al. 2015), *Scytalopus* has not extensively invaded the tropical lowlands except for parts of the Brazilian Atlantic Forest nor undertaken back-colonization to southern South America. In fact, ecological niches as evidenced by elevational distributions exhibit phylogenetic signal in *Scytalopus* (Cadena and Céspedes 2019), with major lineages following distinct elevational zones, such as the *magellanicus* group in the cool treeline/paramo zone, the *latebricola* group in upper cloud forest, the *vicinior* group mainly in lower cloud forests in the western cordilleras, and the *atratus* group in sub-Andean forests and foreland ridges.

A salient feature of tropical montane avifaunas is that species have restricted elevational ranges and replace each other in different elevational zones (Terborgh 1977; Diamond 1973). Species of *Scytalopus* are no exception, showing remarkable turnover along some elevational transects, particularly in areas of high regional species richness. For example, on the western slope of the Andes of Colombia or on the eastern slope of the Peruvian Andes one may find 4-6 *Scytalopus* species with abutting elevational ranges (Stiles et al. 2017; Hosner et al. 2013). Given their poor dispersal abilities which presumably restrict gene flow along elevational gradients and preclude the crossing of deep – and often dry– valleys or cool high-elevation passes, *Scytalopus* tapaculos would appear to be prime candidates for parapatric speciation on mountain slopes (Patton and Smith 1992). However, we found no definitive cases in which sister species replace each other in adjacent elevational zones on the same slope. Instead, sister species typically occur in allopatry and close relatives have similar elevational distributions (Cadena and Céspedes 2019). This result is consistent with work on other montane birds (Caro et al. 2013; García-Moreno and Fjeldså 2000; Moyle et al. 2017) and other vertebrates (Patton and Smith 1992), suggesting that speciation in tropical mountains largely occurs in allopatry within elevational zones and not as a consequence of adaptation to conditions changing with elevation (Cadena et al. 2012). Therefore, coexistence of species along elevational gradients reflects secondary contact following allopatric divergence, implying that geographic ranges of species must have been dynamic over evolutionary time regardless of whether species are dispersal-limited at present (Cadena et al. 2019; Cadena and Céspedes 2019). In this regard, a contrast exists between the radiations of *Scytalopus* in the Andes and in eastern Brazil; whereas in areas of the central and northern Andes elevational replacements of multiple species are commonplace, species rarely segregate by elevation in Brazil. While this may simply reflect that elevational gradients are not as extensive in the eastern Brazilian highlands, it appears that diversification of *Scytalopus* in this region has been largely allopatric, with few cases of lineages reaching secondary sympatry. In cases where species do come into geographic contact, they tend to segregate by habitat more so than by elevation (Maurício 2005; Maurício et al. 2014). Additional analyses of the geographic mode of speciation and elevational replacements in *Scytalopus* are discussed elsewhere (Cadena and Céspedes 2019).

### Diversification dynamics: *Scytalopus* as great speciators

Our estimates of rates of species accumulation in *Scytalopus* obviously differed when employing different data sets, being higher with the taxonomically liberal 83-tip dataset than with the conservative 48-tip data set. True rates are likely somewhere in between, but nonetheless the speed with which lineages of *Scytalopus* accumulated, particularly during the Pliocene, appears rather unusual compared with many other avian groups. For instance, the speciation rates we estimated across the genus using BAMM exceeded those of main clades of hummingbirds (McGuire et al. 2014) and of a clade showing exceptional diversification in the large Neotropical family Furnariidae (i.e. Cranioleuca spinetails; Derryberry et al. 2011; Seeholzer et al. 2017). Rates observed in *Scytalopus* were comparable to those of iconic examples of rapid radiation in tanagers (Thraupidae), including Darwin’s finches and *Sporophila* capuchinos (Burns et al. 2014), as well as to those of some Old World clades like African weavers, monarch flycatchers, *Acrocephalus* warblers, *Aplonis* starlings and *Zosterops* white-eyes (Moyle et al. 2009; J. Fjeldså et al., unpubl. data). In contrast to several of those examples, however, *Scytalopus* species have not diverged extensively in traits related to feeding ecology, habitat, or plumage coloration, suggesting that their rapid diversification was not driven by adaptive processes or sexual selection acting on plumage. Diversification dynamics in *Scytalopus* resemble patterns observed in widespread passerine bird familes in which repeated allopatric speciation resulted in multiple forms with conserved morphology, and contrast with those of lineages diversifying in smaller geographical areas in which diversity was built up thanks to extensive eco-morphological differentiation (Kennedy et al. 2018). In keeping with our findings, work on various organisms suggests that rates of diversification of clades with little phenotypic disparity (Kozak et al. 2006; Valente et al. 2010; Rowe et al. 2011; Moyle et al. 2009) may be on par with those of some textbook examples of rapid adaptive radiation. We hypothesize that extensive non-adaptive diversification in *Scytalopus* resulted from an interaction between traits intrinsic to these organisms such as low dispersal capability making them speciation-prone and the history and topography of the region they inhabit, which provided ample geographical opportunities for allopatric speciation.

Tapaculos are, in general, birds of the lower strata of their forest or thicket habitat, which rarely leave vegetation cover and fly only sporadically in short, fluttering flights. Members of *Scytalopus, Myornis* and *Eugralla* have apparently gone further in the direction of flightlessness, as they have a reduced keel and their clavicles do not fuse into a furcula but consist of two slender spikes (Maurício et al. 2008; Feduccia 1999). The number of tail feathers varies among individuals of a single species of *Scytalopus* (8-14) and some have asymmetric tails, suggesting that the development of the tail is not functionally important; this also applies to their body plumage because moults are remarkably irregular, often with assymetries. *Scytalopus* are ground-living to an extent hardly observed in other passerine groups, tending to stay in the dark, deepest stratum of the vegetation, where they even enter cavities below tree roots or fallen logs and underneath boulders (Krabbe and Schulenberg 1997). Sedentariness associated with near-flightlessness may in some cases lead to increased risk of extinction (Steadman and Martin 2003; Sandel et al. 2011), but limited dispersal may also predispose lineages to become speciation prone a result of greater opportunities for allopatric differentiation (Claramunt et al. 2012; Smith et al. 2014; Greenberg and Mooers 2017). We suggest that because of their limited potential to disperse from one highland region to another, *Scytalopus* populations readily become geographically isolated. Because these small birds are able to build up dense local populations, budding isolates persist over time thus enabling divergence and resulting in the accumulation of diversity (Dynesius and Jansson 2014; Rabosky 2016).

Our reasoning that populations of *Scytalopus* are prone to become isolated and diverge because individuals exhibit poor dispersal abilities may appear counter to the observation that the genus ranges widely in the Neotropics, having managed to expand its distribution extensively across geography. However, the same is true of the diversification of several other birds considered “great speciators”, whose ancestors were able to colonize multiple isolated areas yet subsequently diverged in isolation (e.g. in island archipelagos in the Pacific; Diamond and Mayr 1976). Whether such patterns reflect shifting dispersal abilities over time or rather the influence of changes in the enviroment promoting cycles of range expansions and contractions, the interplay between dispersal and subsequent divergence across geographic barriers appears to have been crucial for the accumulation of avian diversity in both lowland (Smith et al. 2014) and montane (reviewed by Cadena et al. 2019) Neotropical habitats.

We found that climatic dynamics of the Pleistocene leading to fragmentation and reconnection of vegetation belts were unlikely major drivers of the rapid diversification and accumulation of diversity observed in *Scytalopus* because the majority of speciation events occurred prior to the onset of climatic fluctuations of the Late Quaternary. In contrast, LTT plots suggested a downturn in accumulation of lineages towards the present and BAMM analyses revealed an overall decline in speciation rates since the origin of *Scytalopus* in the Miocene. Such patterns of species accumulation over time are consistent with the hypothesis that opportunity for diversification in *Scytalopus* was higher earlier on in the history of the clade and subsequently declined. Declining diversification rates are often interpreted as evidence of ecological limits to diversification (Rabosky 2009), such that niche filling precludes the origin and persistence of new lineages owing to competition. Because species of *Scytalopus* rarely coexist locally perhaps due to mutual exclusion associated with their conserved ecology, ecological limits may partly explain the diversification dynamics we observed (Price et al. 2014; Rabosky and Hurlbert 2015); this hypothesis awaits direct tests (Machac et al. 2013). In addition, extensive diversification in *Scytalopus* coincided with periods of substantial tectonic activity resulting in mountain uplift, particularly in the Northern Andes (Gregory-Wodzicki 2000). Because mountain uplift likely drove diversification by promoting geographic isolation of populations and the origin of novel environments (e.g. páramos) occupied by new lineages (Antonelli et al. 2018), declining rates of diversification over time in *Scytalopus* may also reflect reduced opportunities for diversification as mountains reached their modern elevations in the Late Pliocene (Gregory-Wodzicki 2000). Another likely explanation for declining diversification rates in *Scytalopus* is reduced potential for repeated cycles of allopatric speciation owing to infrequent range expansions (Moen and Morlon 2014). Finally, given incomplete knowledge of species diversity in *Syctalopus* (see below), we cannot rule out the possibility that declining rates of species accumulation towards the tip of the tree may be partly artifactual, resulting from incomplete sampling (Cusimano and Renner 2010). However, we observed similar overall patterns when analyzing data only for well-established species and when employing a more liberal taxonomic scheme in which we considered several lineages potentially representing undescribed species.

### Phylogeny, population structure and implications for species level-taxonomy

The results of our analyses are informative for species-level taxonomy by highlighting cases of non-monophyly of species and by flagging distinct lineages within currently accepted species. We did not employ coalescent approaches to formally identify lineages which may represent presumptive species (e.g. Reid and Carstens 2012) because we lacked population-level sampling across species and geography. Nonetheless, genealogical patterns in gene trees and genetic distances suggest that several groups merit additional work to clarify their true diversity and species limits. A noteworthy case was that of *S. atratus*, which we found is paraphyletic with respect to the vocally distinct *S. sanctaemartae* and *S. bolivianus.* Furthermore, genetic differentiation among populations referred to *S. atratus* is substantial, reaching 10.7% sequence divergence in mtDNA (see long branches in Figures 3 and 4). In combination with seemingly marked vocal variation (Schulenberg et al. 2007), our data suggest that *S. atratus* likely comprises multiple species. That *S. atratus* may comprise more than one species is, however, not entirely unexpected given geographic variation in plumage, with several named subspecies in the group. A perhaps more striking case is that of *S. parvirostris*, which despite being monotypic turned out to be one of the main groups in the Tropical Andes clade, consisting of several genetically distinct populations seemingly isolated by geographic barriers known to structure populations of other birds in the Andes of Peru and Bolivia (Valderrama et al. 2014; Gutiérrez-Pinto et al. 2012; Cadena et al. 2007). As with *S. atratus*, vocal differentiation among populations of *S. parviroristris* has been noted (D. F. Lane, unpubl. data; Krabbe and Schulenberg 1997); detailed analyses of song variation across geography are necessary to adequately delimit species in both groups.

We found that *S. latrans intermedius* is sister either to *S. macropus* (species-tree analysis of nuclear data, mtDNA) or to a clade formed by *S. macropus* and *S. gettyae* (concatenated nuclear data). In turn, other subspecies of *S. latrans* formed a clade with several other taxa (see below). These findings are consistent with vocal variation because the voices of *S. l. intermedius* and *S. macropus* are similar and different from those of *S. l. latrans* (Schulenberg et al. 2007). Moreover, in playback experiments individuals of *S. l. latrans* from southern Ecuador did not respond to songs of *S. l. intermedius* (Freeman and Montgomery 2017). The existing evidence thus indicates that *intermedius* deserves species rank. In turn, the remaining populations of *S. latrans* (i.e. excluding *intermedius*), were scattered within a clade also including *S. micropterus, S. femoralis, S. unicolor, S, gettyae, S. acutirostris* and Peruvian populations which may represent distinct species (i.e. specimens from Lambayeque, specimens referred to *femoralis* from Amazonas and San Martín). The apparent paraphyly of *S. latrans* is possibly a result of incomplete lineage sorting resulting from recent divergence among rapidly radiating species as discussed below. However, the pattern might also represent evidence supporting the hypothesis that this taxon comprises more than one species, as suggested by vocal variation (N. K. Krabbe, unpubl. data, recordings deposited in the Macaulay Library and XenoCanto). For example, *S. latrans subcinereus* and the eastern form of *S. latrans* occurring in Ecuador differ vocally and appear to maintain such differences where they come into contact (e.g. in El Arenal, Ecuador). Potential contact zones between *S. l. subcinereus* and northwestern *S. latrans* in Cañar or Azuay, Ecuador, remain unstudied. Distinct vocal differences between populations of *S. latrans* in the west and east of Colombia and Ecuador are congruent with our finding that they differ genetically, and also the population in north-eastern Colombia and adjacent Venezuela is vocally and genetically distinct. Evidently, species limits within the *S. latrans* complex require additional study.

Intriguingly, our finding that birds referred to as *S. altirostris* represent two distinct mitochondrial and nuclear lineages not closely allied to each other suggested a cryptic species may be involved. Indeed, upon examining recordings of vocalizations we realized there are two song types attributed to *S. altirostris* as well. Usually only one song type occurs at a site, but sometimes the two co-occur. The geographic distribution of song types is generally similar, with one of them appearing to extend farther north in Amazonas, Peru. Because vocal types are locally syntopic and both might occur at the type locality of *S. altirostris* (Atuén, Amazonas; Zimmer 1939), resolving this taxonomic riddle and appropriately dealing with nomenclatural issues will require obtaining sequence data from the type specimen to determine to which population it corresponds. Whichever one of the populations is not *S. altirostris* is most likely a species awaiting description.

Another example of a currently accepted species that may not be monophyletic is *S. acutirostris*, which we found consists of two lineages seemingly not sister to each other. The case of *S. acutirostris* is especially complicated because the type locality of this taxon is uncertain; the original description specified only “Peru” (Tschudi 1844), yet based on the collector’s travels, Cory and Hellymayr (1924) later restricted the locality to Maraynoc, Junín. Krabbe and Schulenberg (1997) indicated that two vocally distinct populations of small, dark-bodied *Scytalopus* occur with elevational segregation in forests in central Peru. These authors tentatively applied the name *S. acutirostris* to the higher-elevation form, whereas the lower-elevation form was treated as a member of the *S. parvirostris* complex. However, because these taxa are impossible to tell apart based on plumage and morphometrics, uncertainty persists regarding to which population does Tschudi’s type specimen corresponds. Our finding of two distinct clades of *S. acutirostris* adds yet another layer of complexity to the situation. Vocally, these two clades differ slightly, but noticeably from each other, while a population further north (Amazonas, La Libertad) for which no sequence is available, is vocally more distinct (N. K. Krabbe, unpubl. data, recordings deposited in the Macaulay Library and XenoCanto). As with *S. altirostris,* analyzing DNA sequences from the type specimen of *S. acutirostris* seems like the only possible way to clarify this taxonomic and nomenclatural riddle. Otherwise, it might be advisable to declare *S. acutirostris* a *nomen dubium*, and describe whichever many new taxa are necessary once geographic variation in vocalizations and genetic structure are better understood.

Additional cases requiring attention to clarify species limits include *S. speluncae,* which as shown by earlier work includes a diversity of lineages (Pulido-Santacruz et al. 2016), and *S. meridanus* in which two distinct and deeply divergent lineages showing small differences in introductions to songs (Donegan and Avendaño-C. 2008) occur at the extremes of the Cordillera de Mérida of Venezuela. The population to the east likely corresponds to form *fuscicauda*, which is currently considered a synonym of *meridanus*, but given our results it may represent a valid taxon. In other cases (e.g. *S. griseicollis*), the existence of distinct lineages had already been noted yet based on additional evidence researchers concluded populations are best considered conspecific (Avendaño and Donegan 2015; Donegan and Avendaño-C. 2008).

In contrast to the cases where marked genetic structure existed within currently accepted species, a few well-established taxonomic species were not recovered as highly divergent mtDNA lineages. In particular, *S. micropterus, S. femoralis, S. unicolor, S, gettyae, S. acutirostris, S. latrans* cf. *latrans, S. latrans subcinerus,* and an undescribed taxon from Lambeyeque and Cajamarca, Peru, are all very similar in mtDNA. These taxa, however, are diagnosable morphologically and their songs are distinctive (Krabbe and Schulenberg 1997; Hosner et al. 2013). Some of them have parapatric distributions (*S. latrans* with *S. micropterus, S. l. subcinereus* with the Lambayeque-Cajamarca form, S. *gettyae* with *S. acutirostris*) and maintain their vocal integrity, which we believe is strong evidence of reproductive isolation. Therefore, the lack of marked mtDNA divergence among taxa in this group is best explained as evidence of recent and rapid speciation, perhaps in the face of gene flow owing to divergence along elevational gradients (Cadena and Céspedes 2019).

We close by noting that although much remains to be clarified about species diversity and species limits in *Scytalopus*, the task no longer seems as daunting as it once appeared. An example of substantial taxonomic progress is that of the *latebricola*-*vicinior* group, in which several populations were elevated to species status based on their vocalizations and mutiple species and subspecies have been described over the past two decades (Krabbe & Schulenberg 1997, Cuervo et al. 2005, Krabbe et al. 2005, Donegan & Avendaño 2008, Donegan et al. 2013, Avendaño & Donegan 2015, Avendaño et al. 2015, Stiles et al. 2017). Although vocal variation (Krabbe et al. 2006; N. Krabbe unpubl. data) and mtDNA differentiation (this study) suggest that *S. spillmanni* and *S. meridanus* may comprise more than one species, and although fieldwork in previously unexplored regions may lead to the discovery of new taxa, we believe that the taxonomic revision of the *latebricola*-*vicinior* group is now largely complete. The detailed studies of Brazilian populations suggest that understanding of lineage diversity in that region is comprehensive, although taxonomic revision in the *S. speluncae* group is still required. Particular attention should now be directed to groups in which difficulties in species delimitation persist including *S. atratus, S. parvirostris*, and the Southern Andes clade, in which our results showing marked genetic divergence among named and unnamed taxa largely confirm species limits suggested by vocal differentiation (N. Krabbe et al. unpubl. data). The molecular data sets and phylogenetic hypotheses we developed for this study represent a baseline framework to guide future taxonomic analyses.

## Supporting information

Supplementary Table 1 - specimens included in mtDNA analyses

Supplementary Table 2 - specimens included in nuclear DNA analyses

## Acknowledgements

A study of this sort would have been impossible without support from multiple natural history collections over many years. We thank the following institutions for providing tissue samples from specimens under their care: Academy of Natural Sciences of Drexler University (N. Rice), American Museum of Natural History (P. Sweet and J. Cracraft), Coleção de Ornitologia do Museu de Ciências e Tecnologia da Pontifícia Universidade Católica do Rio Grande do Sul (C. Fontana), Colección Ornitológica Phelps (J. Pérez-Emán and M. Lentino), Harvard University Museum of Comparative Zoology (S. V. Edwards and J. Trimble), Instituto Alexander von Humboldt (J. D. Palacio, D. López and M. Álvarez), Instituto de Ciencias Naturales de la Universidad Nacional de Colombia (F. G. Stiles), Marjorie Barrick Museum (J. Klicka), Museo de Ciencias Naturales La Salle (M. Salcedo), Museu Paraense Emílio Goeldi (A. Aleixo), The Field Museum (D. Willard and J. Bates), The University of Kansas Natural History Museum (A. T. Peterson, R. Moyle, and M. Robbins), United States National Museum of Natural History (G. Graves and R. T. Chesser), and University of New Mexico Museum of Southwestern Biology (C. Witt and A. Johnson). We are especially grateful to the many collectors who contributed specimens of these difficult birds to museum collections over decades. For collaboration at various stages of this project we thank our colleagues working on Brazilian *Scytalopus* H. Mata, P. Pulido, G. Maurício, M. Bornschein, R. Belmonte-Lópes, and S. Bonatto. Financial support was provided through grants by the Chapman Fund of the American Museum of Natural History and the Facultad de Ciencias at Universidad de los Andes to CDC; the National Science Foundation – NSF to RTB (DEB-1146265, DEB-0910285) and EPD (DEB-1146423); the São Paulo Research Foundation – FAPESP to GAB and LFS (2012-23852-0); the Brazilian National Council for Scientific and Technological Development – CNPq to GAB and LFS (457974-2014-1); and the Society of Systematic Biologists, the Alexander Wetmore Memorial Research Fund of the American Ornithologists’ Union, and the Louis Agassiz Fuertes Award of the Wilson Ornithological Society to AMC. Eugenio Valderrama, Jaqueline Battilana, Renata Beco, Leandro Neves and Rapid Genomics provided valuable assistance with laboratory work. Genomic analyses were run on the Odyssey cluster supported by the FAS Division of Science, Research Computing Group at Harvard University.

---

Supplementary Figures (shown below and in the following pages). Geographic distribution of species in each of the major clades of *Scytalopus* (polygons obtained from BirdLife International) and our sampling for our molecular phylogenetic analyses (symbols).

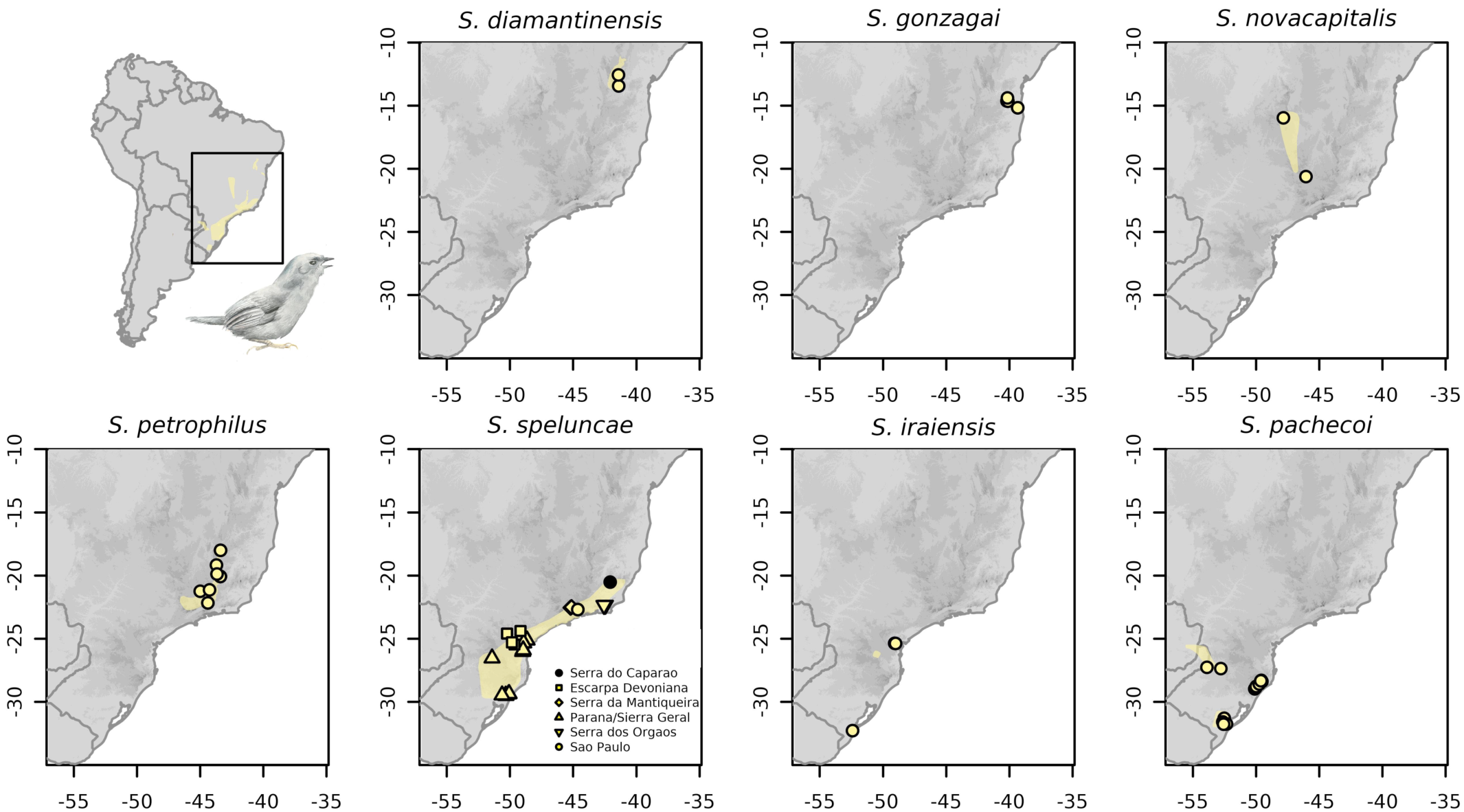
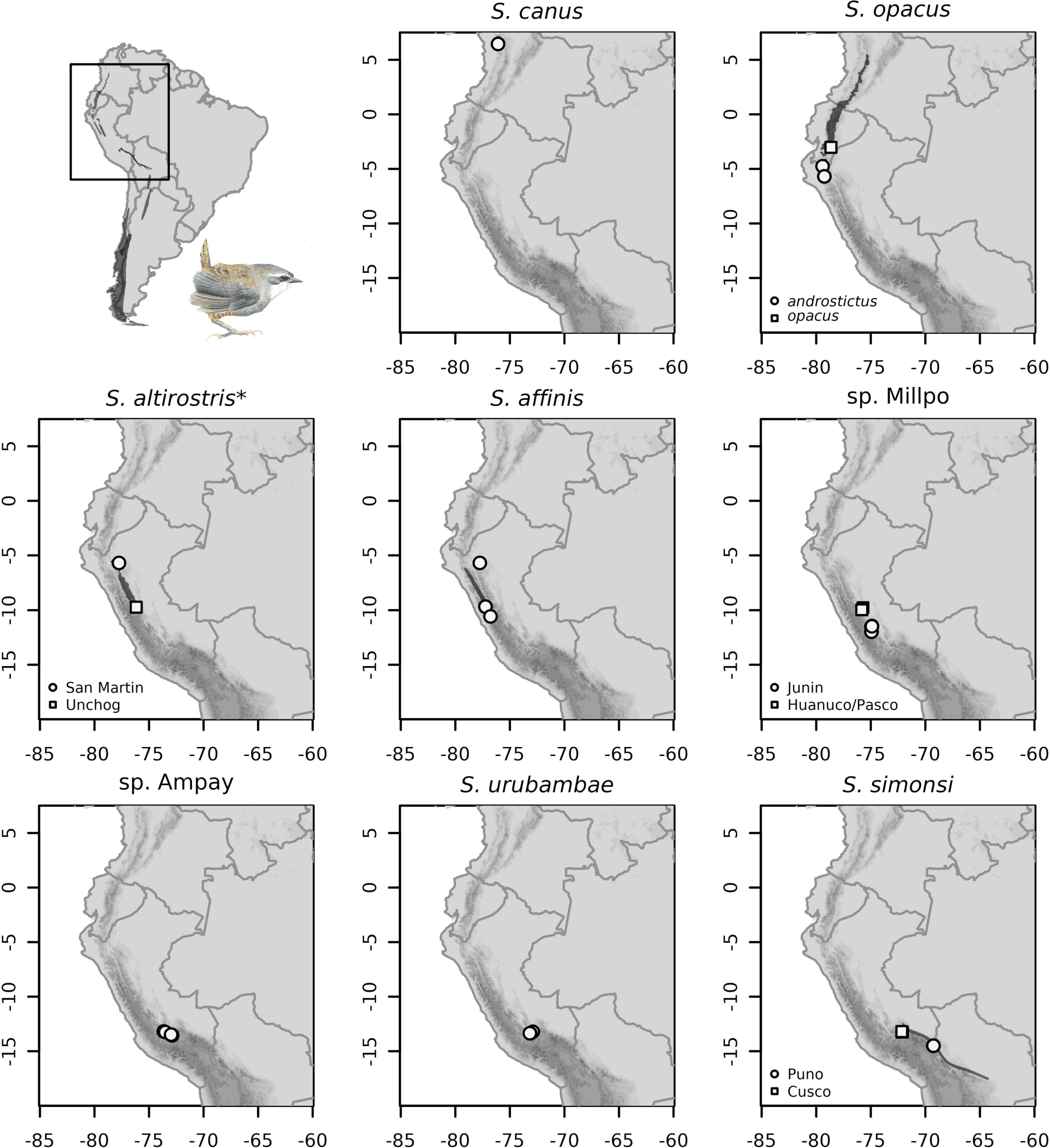
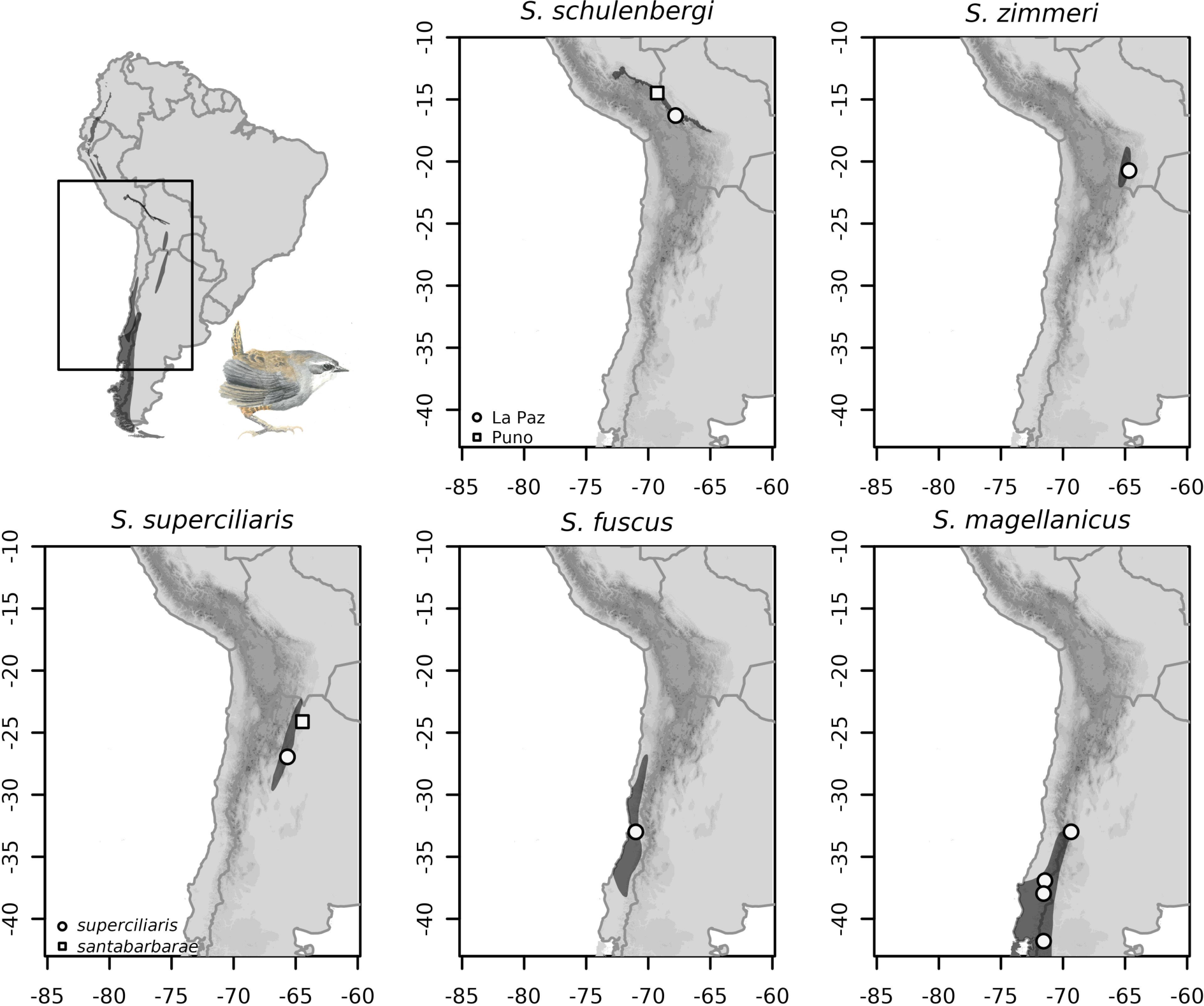
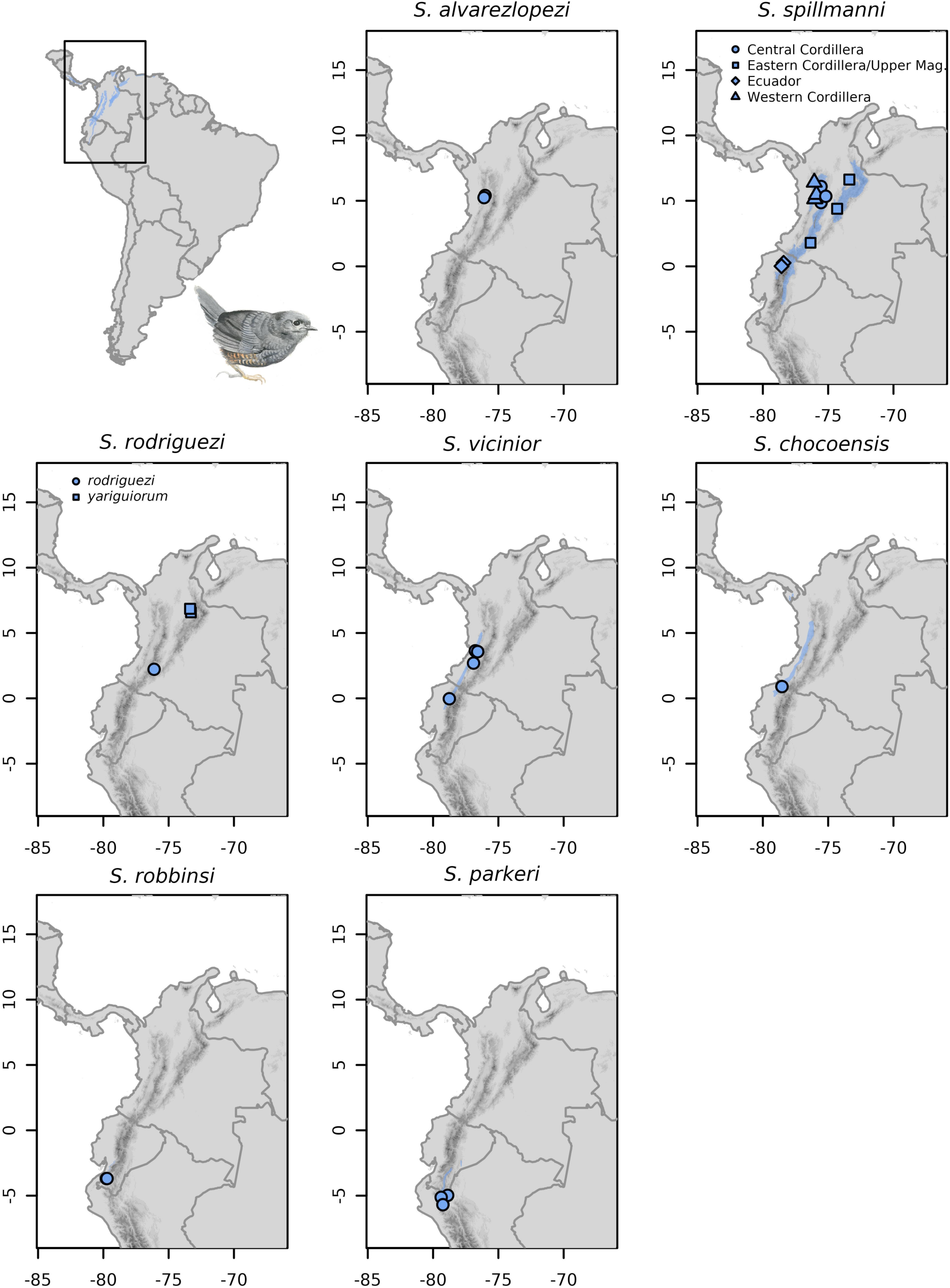
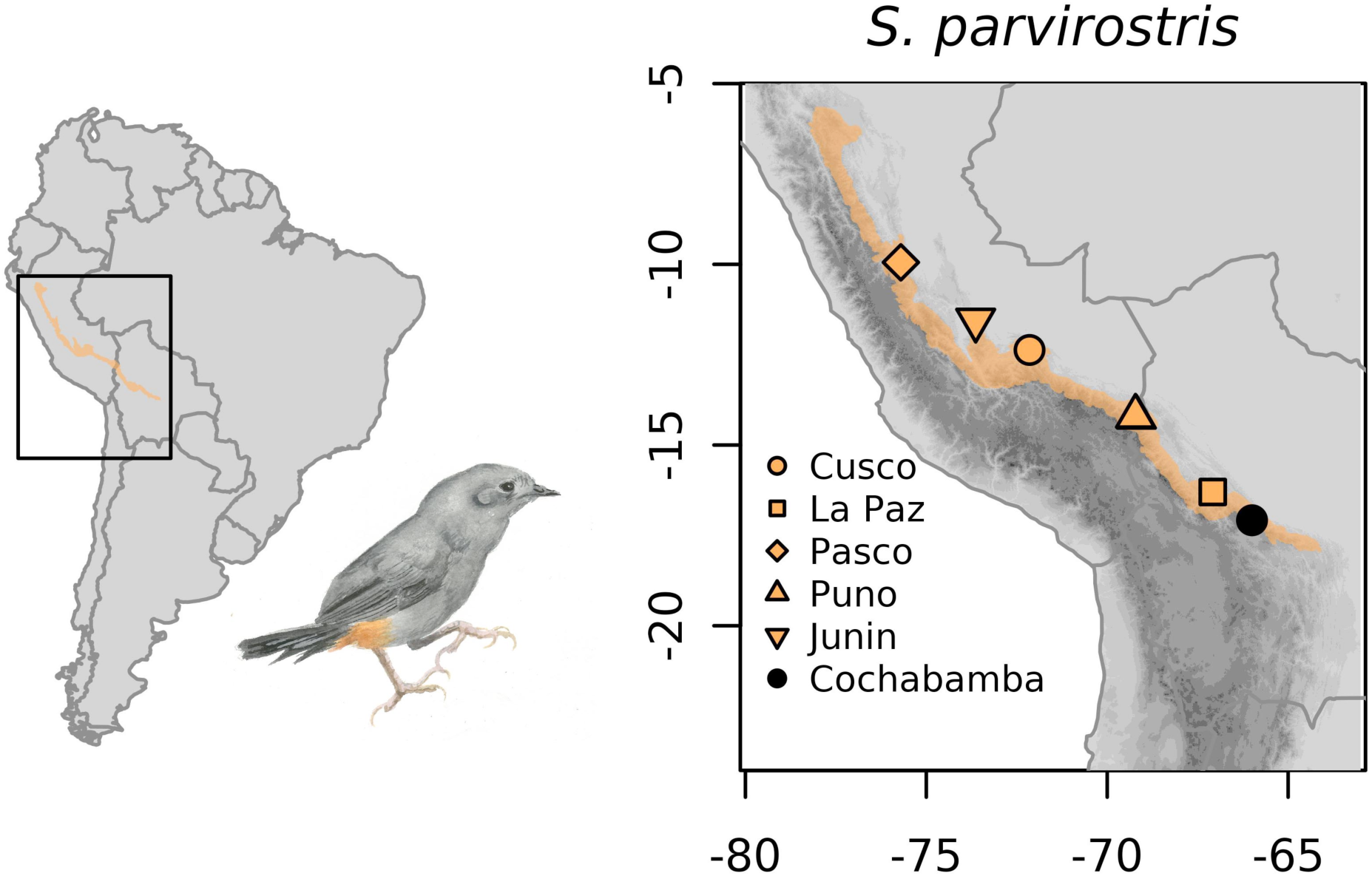
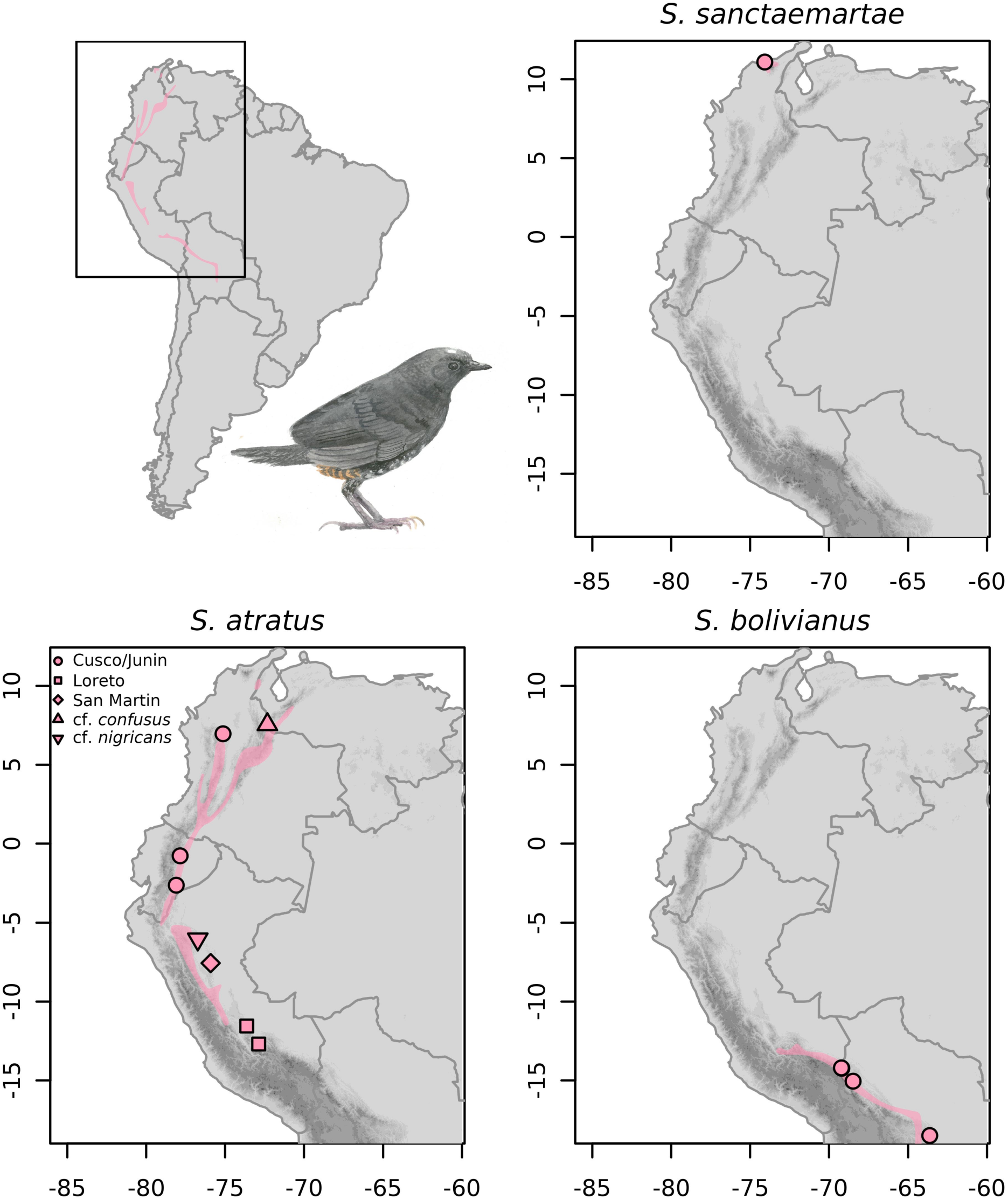
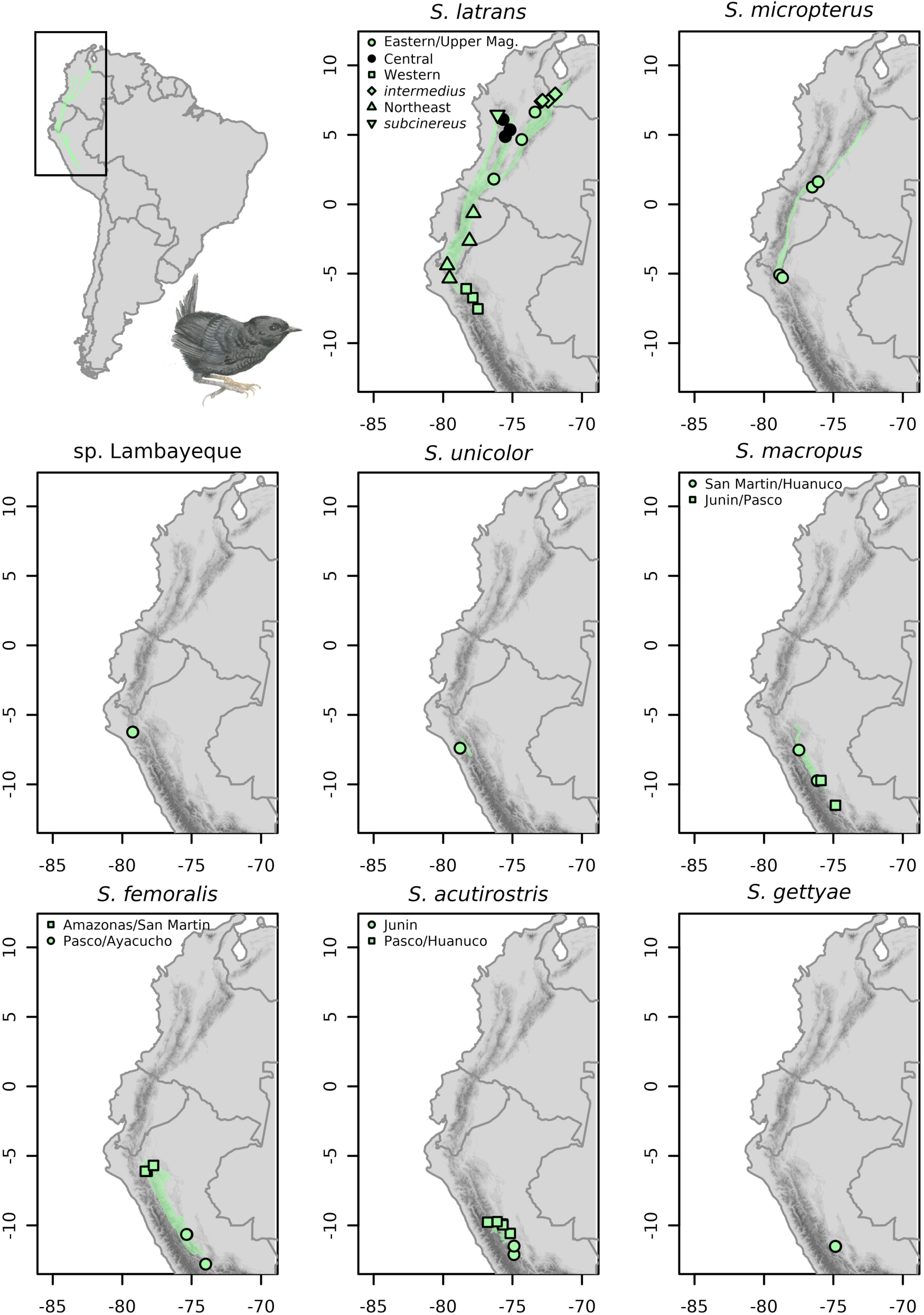
A.Brazilian Clade. B.Southern Andes Clade. C.Southern Andes Clade (continued). D.Tropical Andes Clade: the *latebricola*/*vicinior* group. E.Tropical Andes Clade: the *latebricola*/*vicinior* group (continued). F.Tropical Andes clade: the *Scytalopus parvirostris* complex. G.Tropical Andes clade: the *Scytalopus atratus* complex. H.Tropical Andes clade: the *Scytalopus femoralis-latrans* group.

